# Altered Nucleoprotein Binding to Influenza Virus RNA Impacts Packaging Efficiency and Replication

**DOI:** 10.1101/648261

**Authors:** Valerie Le Sage, Jack P. Kanarek, Eric Nturibi, Adalena V. Nanni, Dan J. Snyder, Vaughn S. Cooper, Seema S. Lakdawala, Nara Lee

## Abstract

The genome of Influenza A viruses consists of eight negative-sense RNA segments that are bound by viral nucleoprotein (NP). We recently showed that NP binding is not uniform along the segments but exhibits regions of enrichment as well as depletion. Furthermore, genome-wide NP binding profiles are distinct even in strains with high sequence similarity, such as the two H1N1 strains A/WSN/1933 and A/California/07/2009. Here, we performed interstrain segment swapping experiments with segments of either high or low congruency in NP binding, which suggested that a segment with a similar overall NP binding profile preserved replication fitness of the resulting virus. Further sub-segmental swapping experiments demonstrated that NP binding is affected by changes to the underlying nucleotide sequence, as NP peaks can either become lost or appear *de novo* at mutated regions. Unexpectedly, these local nucleotide changes in one segment not only affect NP binding *in cis*, but also impact the genome-wide NP binding profile on other segments in a vRNA sequence-independent manner, suggesting that primary sequence alone is not the sole determinant for NP association to vRNA. Moreover, we observed that sub-segmental mutations that affect NP binding profiles can result in reduced replication fitness, which is caused by defects in vRNA packaging efficiency and an increase in semi-infectious particle production. Taken together, our results indicate that the pattern of NP binding to vRNA is important for efficient virus replication.

**Author Summary:** Each viral RNA (vRNA) segment is bound by the polymerase complex at the 5′ and 3′ ends, while the remainder of the vRNA is coated non-uniformly and non-randomly by nucleoprotein (NP). To explore the constraints of NP binding to vRNA, we used high-throughput sequencing of RNA isolated by crosslinking immunoprecipitation (HITS-CLIP) of mutant H1N1 strains with exchanged vRNA sequences and observed that NP binding can be changed based on vRNA sequence. The most striking observation of our study is that nucleotide changes in one segment can have genome-wide effects on the NP binding profile of other segments. We refer to this phenomenon as the ‘butterfly effect’ of influenza packaging. Our results provide an important context in which to consider future studies regarding influenza packaging and assembly.

## Introduction

The segmented nature of influenza A virus (IAV) genomes poses a logistical challenge for viral replication, as all of the eight negative-sense single-stranded RNA segments must find their way into a budding virion to give rise to an infectious particle [1, 2]. Following nuclear export, viral ribonucleoprotein complexes (vRNP) containing newly synthesized viral RNA (vRNA) assemble on recycling endosomes en route to the plasma membrane for packaging into virions [3, 4]. An accumulating body of evidence suggests that the intracellular pre-assembly process of vRNP trafficking is mediated by RNA-RNA interactions between segments [5–10], which is substantiated by *in vitro* RNA binding studies indicating that multiple sites within vRNA segments form RNA-RNA interactions [8, 11]. Further support for these intersegmental interactions comes from colocalization studies during intracellular viral assembly that showed that segments colocalize with certain other segments preferentially during their transport to the plasma membrane [3, 4].

All eight IAV segments are coated by viral nucleoprotein (NP) molecules, which until recently were thought to cover the entire length of the segments uniformly like ‘beads on a string’ [12–17]. Using HITS-CLIP (high throughput sequencing of RNA isolated by crosslinking and immunoprecipitation), we have previously demonstrated that NP binding to vRNA in virions is not regular but enriched at some regions of the segments while depleted at others, providing evidence for an alternative model of non-uniform NP association to vRNA [18]. The advantage of HITS-CLIP is that, unlike other versions of the CLIP methodology, such as iCLIP or eCLIP [19], it uncovers the entire footprint of vRNA protected by NP rather than focusing on the NP-crosslinked sites on vRNA. Another study utilizing PAR-CLIP (photoactivatable ribonucleoside-enhanced crosslinking and immunoprecipitation) [20], a technically related version of HITS-CLIP that can also identify NP binding sites with nucleotide resolution, reached the same conclusion that NP binding is not pervasive throughout the segments inside infected host cells [21].

We have further shown that the NP binding profiles of strains with a high degree of nucleotide sequence conservation can differ markedly [18]. The A/WSN/1933 (WSN) and A/California/07/2009 (H1N1pdm) strains, both of the H1N1 subtype with an overall sequence homology of 85%, contain NP binding sites that are shared between both strains as well as unique to each strain. Previous *in vitro* binding assays indicated that NP binds RNA in a sequence-independent manner [22, 23], raising the question of how strain-specific NP binding to vRNA is accomplished. We did observe a statistically robust bias for NP binding sites in that they are relatively depleted in uracils and enriched for guanines compared to genome-wide nucleotide content [18]. However, given the vast spread in nucleotide content among all NP peaks in the viral genome, this bias is unlikely to be the sole underlying determining factor of NP recruitment. Moreover, despite the lack of nucleotide selectivity of NP *in vitro*, it cannot be ruled out that accessory proteins *in vivo* ensure specific nucleotide sequences to be recognized and bound by NP. An alternative possibility is that three-dimensional organization of the IAV genome guides NP interaction with vRNA, which would be somewhat comparable to higher-order chromatin structure of eukaryotic DNA genomes contributing to nucleosome packaging [24, 25].

In this study, we examined how NP association impacts virus replication by introducing local mutations to alter the NP binding profile. We unexpectedly observed that local changes in nucleotide sequence can produce global changes in NP binding in a nucleotide sequence-independent manner. Moreover, we observed that alterations in NP binding profiles can affect virus replication kinetics by adversely affecting segment packaging efficiency and increasing the proportion of semi-infectious particles. Taken together, our data indicate an essential contribution of NP binding to vRNA for productive virus assembly.

## Results

### Introducing a segment with a divergent NP binding profile reduces replication fitness

Comparative analysis between the genome-wide NP binding profiles determined by NP HITS-CLIP of the H1N1pdm and WSN strains indicated a varying degree of similarity among segments [18]. For example, the NA segments of both strains show a high Pearson correlation in terms of NP binding (**Figure 1A**), while the NS segments display the lowest Pearson correlation coefficient of all segments (**Figure 1B**). This variability does not reflect nucleotide conservation, as the NA and NS segments are 81% and 85% conserved, respectively. We sought to determine the relationship between NP-vRNA binding and virus replication by exchanging either the NA or NS segment of the H1N1pdm strain with the equivalent segment of the WSN strain to generate two chimeric mutant strains using reverse genetics. Infection of these strains was performed at a MOI of 0.01 and the dynamics of virus production was measured by TCID50 assays at the indicated time points. Multi-cycle infection experiments showed no significant difference in replication between the mutant virus containing the WSN NA segment within the H1N1pdm background (pdm [WSN NA]) and the wildtype H1N1pdm strain, suggesting that segments with similar NP-vRNA binding profiles can efficiently complement virus replication (**Figure 1C**). In contrast, viral titers of the mutant strain containing the WSN NS segment (pdm [WSN NS]), which differs greatly in the overall NP binding profile, were significantly lower than wildtype at 16, 24, and 48 hours post infection (hpi) (**Figure 1C**). This observation suggests that introducing a segment with a more divergent NP-vRNA binding profile can deleteriously affect virus replication.

**Figure 1.**
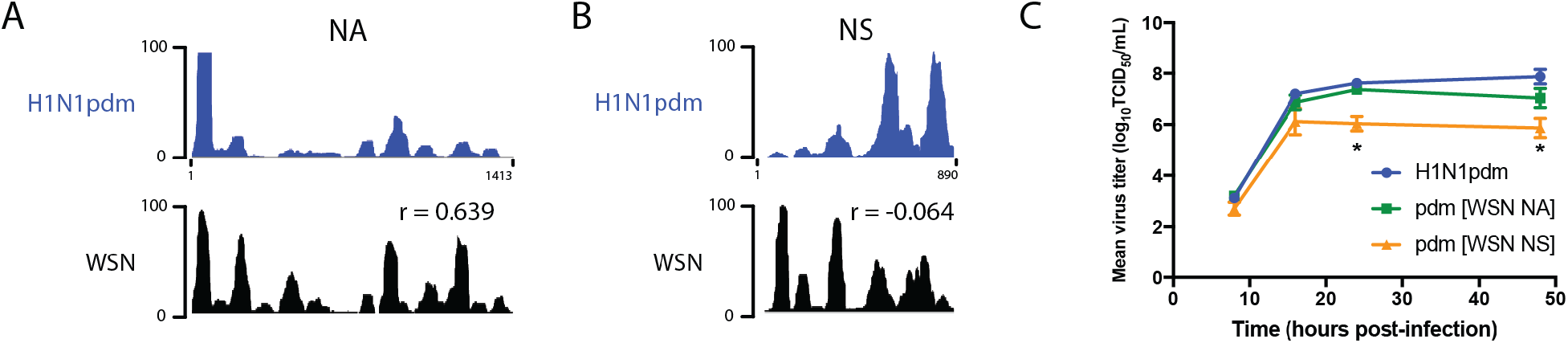
Swapping segments with similar NP binding profiles preserves replication fitness. (**A+B**) The NP binding profiles for the NA and NS segments are shown for A/California/07/2009 (H1N1pdm) and A/WSN/1933 (WSN) strains. Abundance of CLIP reads (y-axis) was normalized against the highest peak in each individual vRNA segment and arbitrarily set to 100. Sequencing tracks and Pearson correlation coefficients (r) between WSN and H1N1pdm segment pairs are taken from Lee et al. [18]. (C) Replication kinetics of the wildtype and two H1N1pdm mutant strains, for which either the NA or NS segment has been exchanged with the WSN equivalent. MDCK cells were infected at a MOI of 0.01, and supernatants were collected at indicated time points to determine virus titers using TCID_50_ assays. Graphs are representative of three independent experiments. Two-way ANOVA analysis was used to determine statistically significant differences (marked by asterisk).

### vRNA sequence influences NP binding

To further examine the relationship between NP-vRNA binding and virus replication, we sought to alter the NP-vRNA binding profile of a single segment within its own viral background and assess the impact on viral lifecycle. The 5′ regions of the NS segments display the most divergent NP binding profiles between the WSN and H1N1pdm strains, even though their nucleotide sequence varies only in 32 out of 220 nucleotides (85% conservation) (**Figure 2A**). This region in the WSN NS gene segment exhibits robust NP binding, while the corresponding region in H1N1pdm is depleted for NP association (**Figure 2B**, red boxes). Therefore, we introduced sub-segmental swapping mutations and generated chimeric NS segments by placing the NP-bound sequence of WSN (nucleotides 50 to 251) into the H1N1pdm background (referred to as strain pdm [WSN-NS 5′]). A reciprocal mutant virus that contains the NP-free region of H1N1pdm in the WSN NS segment (referred to as strain WSN [pdm-NS 5′]) was also generated (**Figure 2B**). These ~200 nucleotides of the 5′ NS vRNA account for the C-terminal regions of the NS1 and NS2 proteins, which are 80% and 90% conserved, respectively, between both strains. These NS mutant strains were rescued and amplified for subsequent HITS-CLIP analyses to identify their NP binding profiles. Introducing the WSN sequence into the H1N1pdm background resulted in the formation of ectopic NP binding sites, as observed in the WSN strain (**Figure 2C**, top panel). This observation demonstrates that the NP binding profile of a given segment is not static but indeed amenable to nucleotide alterations. Similarly, introduction of the H1N1pdm 5′ NS region into the corresponding locus in the WSN NS segment resulted in loss of these NP peaks, which is reminiscent of the NP binding profile of the H1N1pdm strain (**Figure 2C**, bottom panel). Taken together, our results indicate that vRNA sequence can direct NP binding.

**Figure 2.**
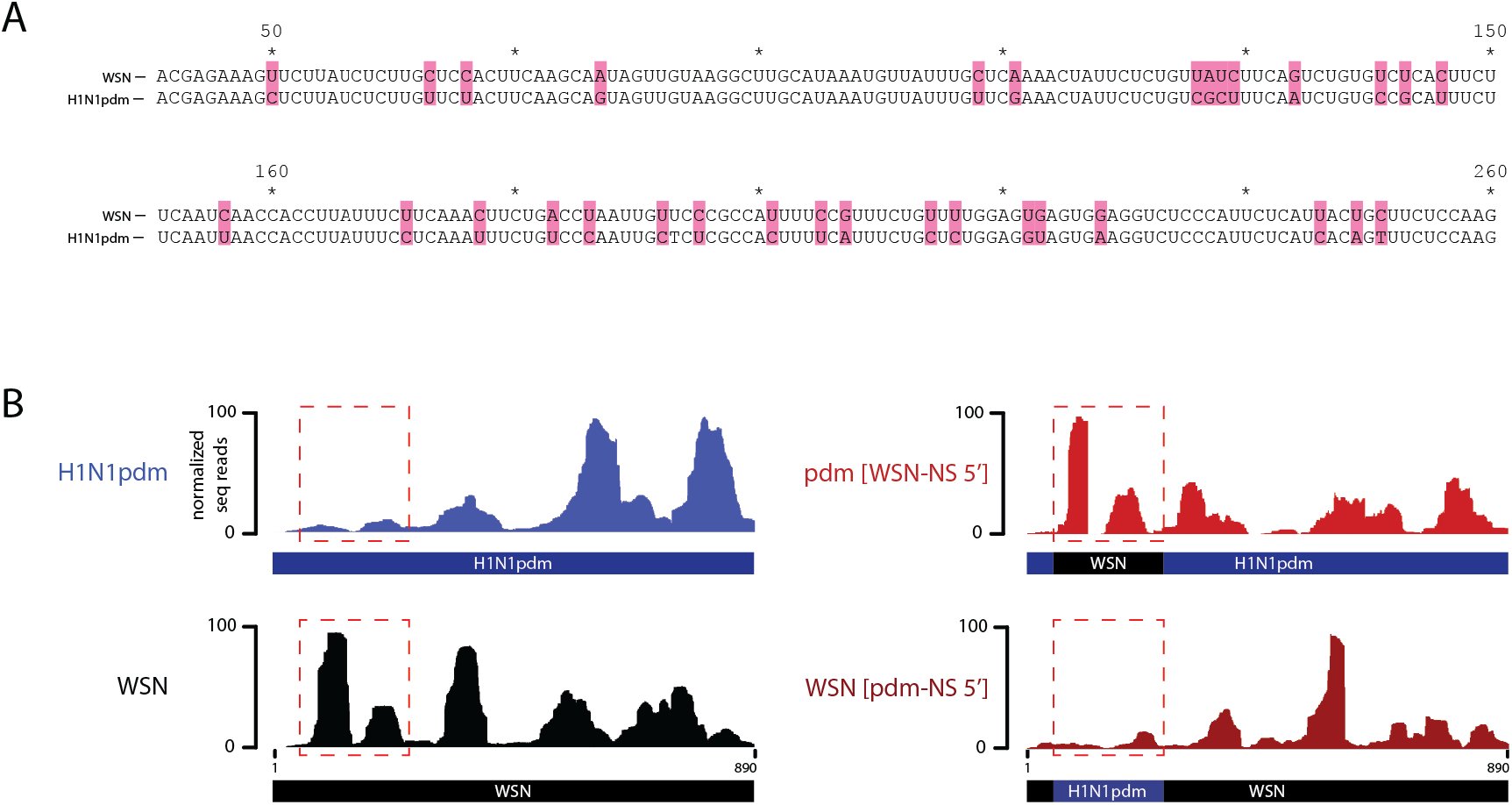
Genomic vRNA mutations can cause alterations in NP binding. (**A**) Sequence alignment of the NS segment 5′ region of the WSN and H1N1pdm strains (red dashed boxes shown in B). Nucleotide differences between the strains are highlighted in pink. Numbers indicate the nucleotide position in the vRNA segment. (**B**) The boxed region of the WSN strain, which contains NP peaks, was swapped with the corresponding NP-free sequence of H1N1pdm, resulting in two chimeric NS segment mutant strains (referred to as pdm [WSN-NS 5′] and WSN [pdm-NS 5′]). NP binding profiles of the NS segments are shown for WSN, H1N1pdm, and the two chimeric strains.

### Local nucleotide changes in vRNA impact NP binding in other segments

Unexpectedly, genome-wide examination of the NP binding profiles of the two NS chimeric mutants revealed that the 5′ region of the NS segment was not the only site that showed a transformed NP binding profile (**Figure 3**). To compare NP peak locations between strains in an unbiased manner, a peak-finding algorithm was used to call specific peaks on each segment and overlap with the peaks of another strain. Our analysis revealed that the majority of NP peaks prevailed in the parental and chimeric mutant strains, yet a number of peaks were detected that are present only in either strain (see **Tables 1+2** for coordinates of all called peaks). In particular, the novel peaks in the NS segment of the chimeric mutant pdm [WSN-NS 5′] strain were noticeably accompanied by loss of NP peaks in the PB2, HA, and M segments, while ectopic peaks emerged in the PB1, PA, and M segments (**Figure 3A**, arrowheads). A similar observation was made when comparing the WSN [pdm-NS 5′] chimeric mutant to its parental WSN strain (**Figure 3B**, arrowheads). Remarkably, all of these alterations in NP-vRNA association occurred in the absence of vRNA nucleotide changes at the respective loci and despite the fact that the primary nucleotide sequence for each of these segments of the chimeric mutants is identical to the parental strains. We verified by re-examining the deep sequencing reads of our HITS-CLIP data that secondary mutations, which may have accumulated during the propagation of the chimeric strains, were indeed absent at the affected loci. Taken together, our findings clearly demonstrate that primary nucleotide sequence *per se* cannot solely account for NP deposition on vRNA. Moreover, this observation is in line with previous *in vitro* studies that indicated that NP binds RNA in a sequence-independent manner [22, 23] and suggests that NP binding specificity is governed by an additional layer of regulation beyond primary nucleotide sequence.

**Figure 3.**
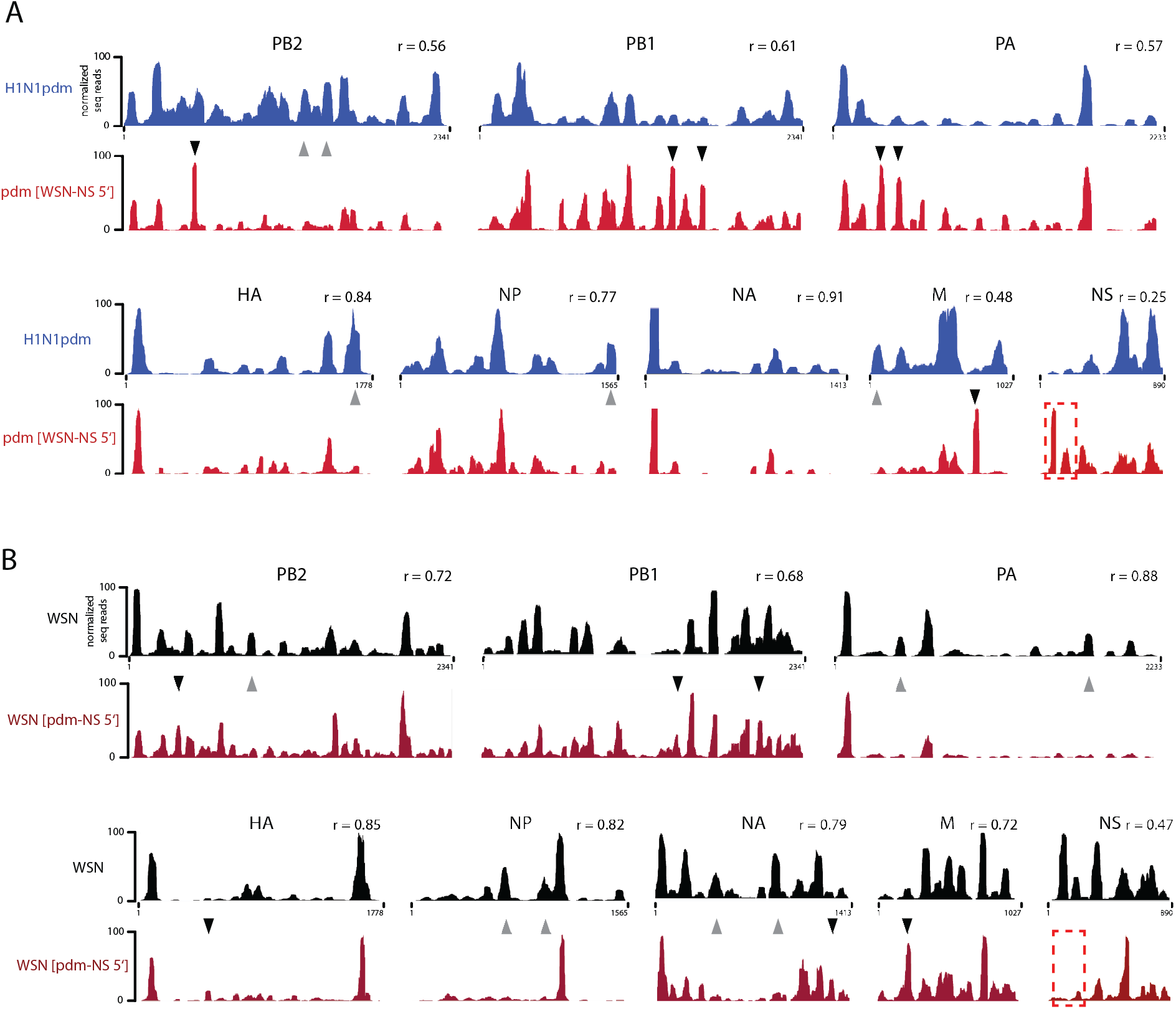
Genome-wide NP binding profile is affected by local mutations independent of underlying vRNA sequence. (**A+B**) Comparison of NP binding profiles determined by HITS-CLIP between wildtype and chimeric mutant H1N1pdm (A) and WSN strains (B). Arrowheads indicate exemplary regions of NP peaks that are noticeably different between wildtype and the chimeric mutant viruses. Tables 1 and 2 list the coordinates of all shared and unique regions. Pearson correlation coefficients (r) between wildtype and chimeric segment pairs are indicated. Red dashed boxes denote the mutated regions. Representative tracks of all eight IAV segments are shown. Note that biological replicates are highly reproducible in their genome-wide NP binding profiles, with Pearson correlation coefficients of >0.7.

**Table 1.**
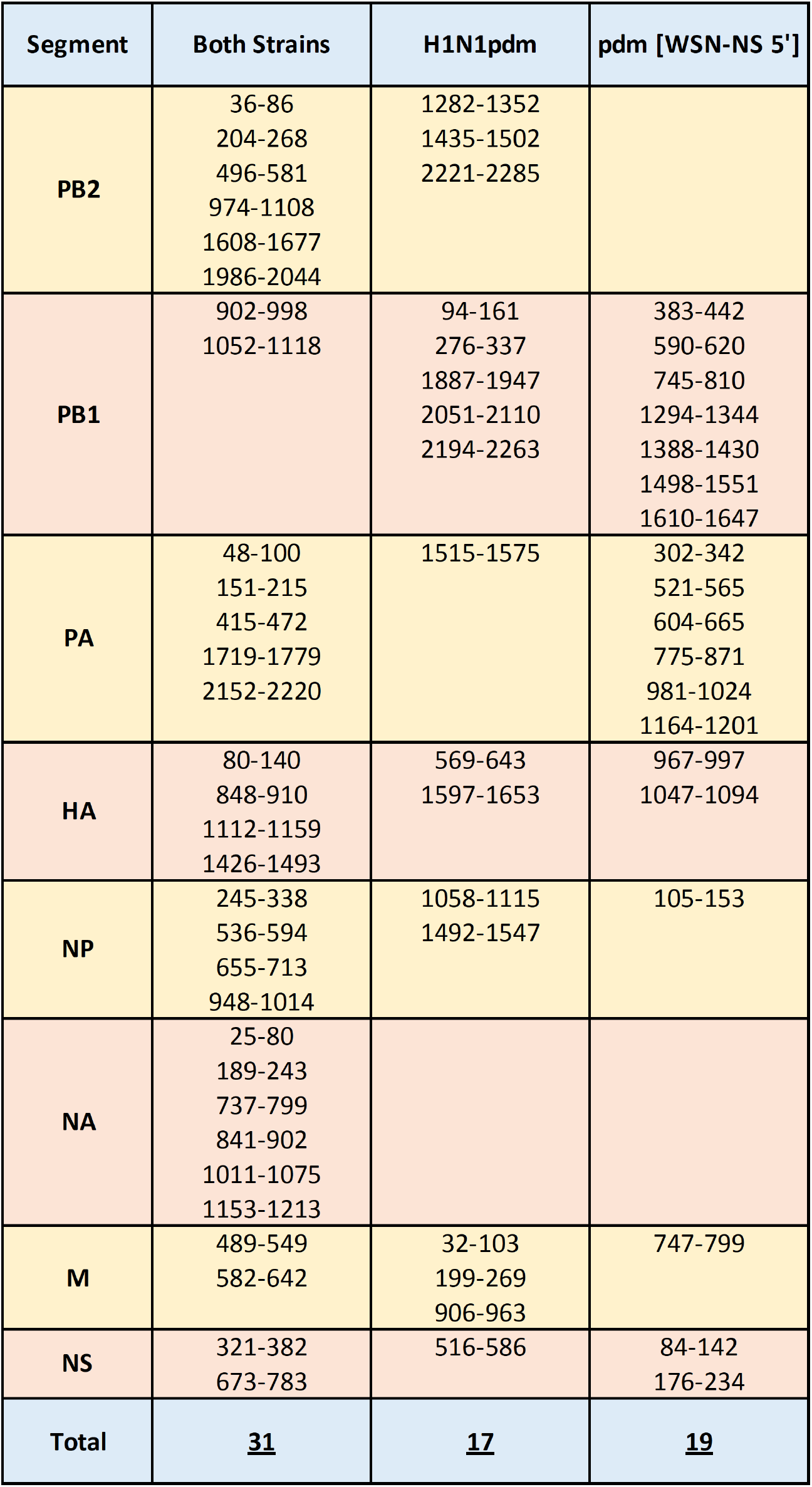
Coordinates of common and unique NP peaks for the wildtype and chimeric NS mutant H1N1 pdm strains (related to **Figure 3A**).

| Segment | Both Strains | H1N1pdm | pdm [WSN-NS 5'] |
| --- | --- | --- | --- |
| **PB2** | 36-86<br>204-268<br>496-581<br>974-1108<br>1608-1677<br>1986-2044 | 1282-1352<br>1435-1502<br>2221-2285 |  |
| **PB1** | 902-998<br>1052-1118 | 94-161<br>276-337<br>1887-1947<br>2051-2110<br>2194-2263 | 383-442<br>590-620<br>745-810<br>1294-1344<br>1388-1430<br>1498-1551<br>1610-1647 |
| **PA** | 48-100<br>151-215<br>415-472<br>1719-1779<br>2152-2220 | 1515-1575 | 302-342<br>521-565<br>604-665<br>775-871<br>981-1024<br>1164-1201 |
| **HA** | 80-140<br>848-910<br>1112-1159<br>1426-1493 | 569-643<br>1597-1653 | 967-997<br>1047-1094 |
| **NP** | 245-338<br>536-594<br>655-713<br>948-1014 | 1058-1115<br>1492-1547 | 105-153 |
| **NA** | 25-80<br>189-243<br>737-799<br>841-902<br>1011-1075<br>1153-1213 |  |  |
| **M** | 489-549<br>582-642 | 32-103<br>199-269<br>906-963 | 747-799 |
| **NS** | 321-382<br>673-783 | 516-586 | 84-142<br>176-234 |
| **Total** | **<u>31</u>** | **<u>17</u>** | **<u>19</u>** |

### Alterations in NP-vRNA binding can affect virus replication by modulating segment packaging efficiency and semi-infectious particle production

To study the impact of NP-vRNA binding changes on the viral lifecycle in the NS 5′ sub-segmental mutants, we next compared the multi-cycle growth kinetics of the chimeric mutant viruses to their parental strains. The pdm [WSN-NS 5′] mutant displayed a comparable replication rate to the parental H1N1pdm strain (**Figure 4A**), while the WSN [pdm-NS 5′] mutant grew to significantly lower titers at 16, 24, and 48 hpi (**Figure 4B**). We reasoned that a decrease in viral replication should also be reflected in segment packaging efficiency into virions. To this end, we conducted competitive plasmid transfection assays between wildtype and chimeric NS segments [26], and generated influenza viruses by transfecting the reverse genetics plasmids containing segments 1-7 together with two distinct NS segments that would compete for incorporation into virions (**Figure 4C**). We performed the competitions for both H1N1pdm and WSN backgrounds by harvesting the rescued viruses and amplifying by RT-PCR a region within the NS segment that spans the swapped locus to distinguish the origin of the NS segment packaged in the progeny virions. The PCR amplicons were then converted into an Illumina-compatible library and deep sequenced to determine the ratio of the incorporated NS segments. In the H1N1pdm background, we competed the wildtype with the chimeric pdm [WSN-NS 5′] segment and observed that the latter did not package less preferentially into progeny virions (40.5% vs. 59.5%; **Figure 4D**). This absence of preference for the wildtype segment may explain why no virus replication defect was observed for the chimeric NS segment. Conversely, competition between the wildtype and chimeric WSN [pdm-NS 5′] segments within the WSN background revealed a significant preference for the wildtype NS segment, as 81.7% of the wildtype segment was found in progeny virions as opposed to 18.3% of the chimeric segment (**Figure 4D**), which provides an explanation for the observed growth defect of the WSN [pdm-NS 5′] mutant strain. Taken together, these data suggest that binding of NP at the 5′ end of WSN NS vRNA is important for its efficient packaging.

**Figure 4.**
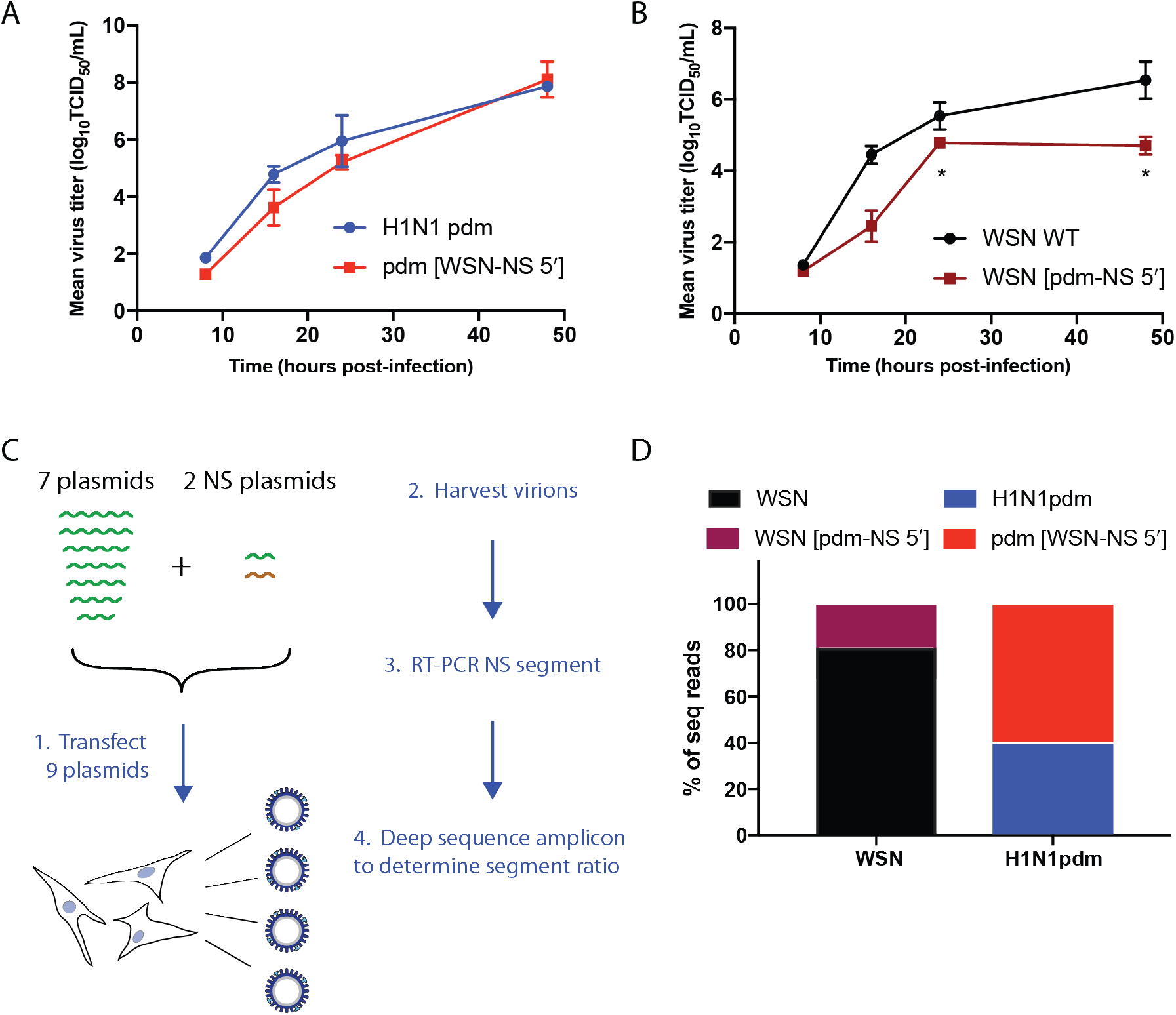
Viral replication and packaging preference of NS chimeric mutants. (**A+B**) Replication kinetics of wildtype and chimeric mutants of H1N1pdm (A) and WSN strains (B). MDCK cells were infected in triplicate at a MOI of 0.01. Supernatants were collected at the indicated time points and virus titers were determined using TCID50 assays. Two-way ANOVA analysis was used to determine statistically significant differences (marked by asterisks). (**C**) Schematic of segment packaging competition assay. Seven plasmids containing either H1N1pdm or WSN segments 1-7 were co-transfected with two distinct NS plasmids as indicated to reverse engineer viruses; the two NS segments compete for packaging into virions. Upon harvesting progeny viruses, vRNA was isolated and subjected to RT-PCR of the NS segment. A primer pair annealing to conserved sequences in both NS variants was used to generate an amplicon, which encompasses the mutated region. Following library preparation, the amplicon was deep sequenced to determine the incorporation ratio of the NS variants into virions. (**D**) Percentage of deep sequence reads of the NS variants incorporated into H1N1pdm or WSN virions after virus rescue. Values are the average of at least two independent biological replicates.

To further examine a potential packaging defect for the chimeric WSN [pdm-NS 5′] mutant, we performed multi-color fluorescence *in situ* hybridization (FISH) to assess segment colocalization during viral infection. We focused on the colocalization of the NS with the M segment, as our previous studies have shown that the intracellular distribution of the M segment is highly correlated with the distribution of the NS segment, and that these two segments cluster together in putative vRNA segment assembly network constructions with machine learning [27]. Host cells infected with either wildtype or chimeric mutant virus for both WSN and H1N1pdm backgrounds were fixed and stained with probes at 8 hpi (**Figure 5A**). The total number of colocated spots within multiple cells was quantified using a previously developed image analysis pipeline [28]; at least five cells were imaged per virus strain on a confocal microscope with fine z-stack to produce a 3D image. A similar number of vRNA spots were identified in each cell, and the proportion of cytoplasmic M or NS foci alone or colocated with each other were measured. A significant decrease in segment colocalization was observed in the WSN [pdm-NS 5′] strain compared to wildtype WSN (**Figure 5B**). Consistently, a higher proportion of cytoplasmic foci contained either M or NS segments alone in the chimeric mutant. On the other hand, we did not detect a difference in segment colocalization between the M and NS segments for the chimeric pdm [WSN-NS 5′] strain (**Figure 5C**), which is in line with the fact that a similar growth rate was observed as for the wildtype virus. Thus, our results indicate that alterations in NP-vRNA association can result in segment colocalization defects during intracellular virus assembly.

**Figure 5.**
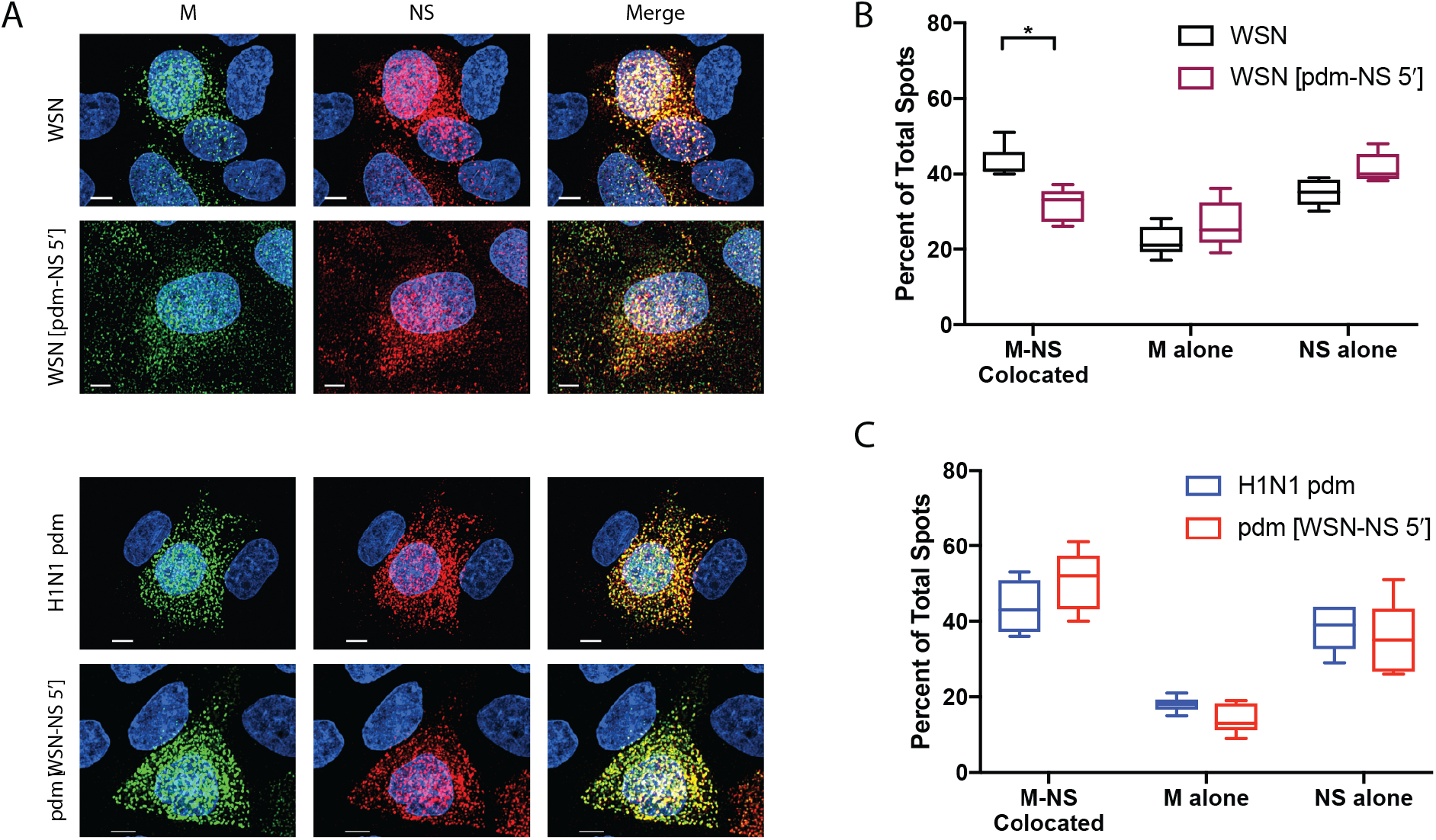
The NS chimeric mutant of the WSN strain shows a segment colocalization defect. (**A**) Representative FISH images of M and NS segments from two independent experiments. MDCK cells were infected with either wildtype or chimeric mutants at a MOI of 3 and then fixed 8 hpi. FISH probes targeting the M vRNA (Alexa 488, green) and NS vRNA (Quasar 570, red) were used. Cell nuclei were stained with DAPI (blue). Scale bars are 5 μm. (**B+C**) Quantification of the cytoplasmic colocalization of M and NS segments for the wildtype WSN and mutant WSN [pdm-NS 5′] strains (B), and H1N1pdm and mutant pdm [WSN-NS 5′] strains (C). Fine confocal stacks were acquired to reconstruct a 3D cell volume. Image analysis on deconvolved stacks included generation of spots for each individual vRNA segment and quantification of colocalization of these spots in the cytoplasm by using DAPI signal to mask the nuclear volume. Five cells were analyzed for each condition. Two-way ANOVA analysis was used to determine statistically significant differences (marked by asterisks).

Previous studies have demonstrated that the production of semi-infectious particles during influenza infection may be a result of inefficient packaging. To determine whether the observed defect in packaging led to an increase in semi-infectious particle development, we compared the total number of particles, quantified by the HA titers, and infectious particles, quantified by plaque titers, of WSN [pdm-NS 5′] to its parental strain. HA titers of WSN [pdm-NS 5′] were comparable to wildtype, whereas PFU per mL of the chimeric virus was reduced 2 to 10-fold (depending on the replicate) (**Figure 6A**). Additionally, we performed qPCR analysis for all eight segments on vRNA extracted from wildtype and mutant WSN [pdm-NS 5′] virions normalized to PFU per mL. The Ct values for the chimeric WSN [pdm-NS 5′] mutant were lower than for the wildtype strain, indicating an overall higher absolute quantity of vRNA in the mutant while the relative abundance between the eight segments within each strain was similar (**Figure 6B**). These data indicate that, in virus preparations with similar infectivity, the WSN [pdm-NS 5′] mutant produced more semi-infectious particles. This conclusion was confirmed by an increase in protein levels of HA, NP, and M1 in the WSN [pdm-NS 5′] strain as compared to wildtype in sample preparations of similar infectious titer (**Figure 6C**). Taken together, these results suggest that the growth defect of the WSN [pdm-NS 5′] mutant is due to a packaging defect, which results in more semi-infectious particles.

**Figure 6.**
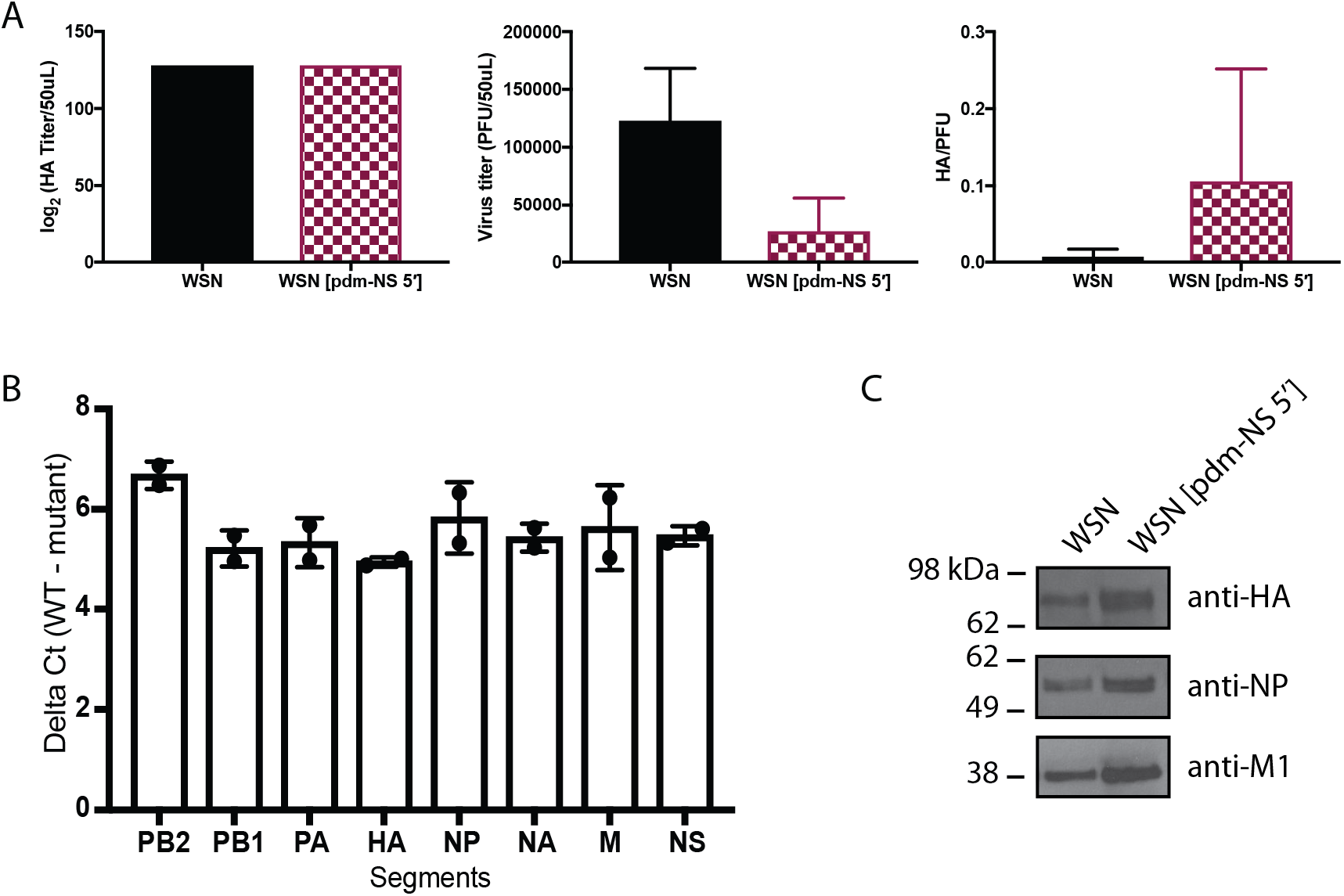
Production of semi-infectious particles is increased in the WSN NS chimeric mutant. (**A**) HA and PFU titers of WSN and NS chimeric virus strains. (**B**) Relative quantification of vRNA segments between wildtype and NS chimeric mutant strains by RT-qPCR. Values are the average of three independent experiments. RNA levels were normalized to PFU titers. (**C**) Supernatants from WSN or NS chimera-infected MDCK cells were collected at 48 hpi. Equal amounts of PFU were concentrated, lysed and the viral proteins separated by SDS-PAGE. Viral proteins were processed for Western blotting and probed for HA, NP and M1. Data shown is a representative of two independent experiments.

## Discussion

We have recently shown that NP binding to vRNA is not pervasive, but restricted to specific regions of the viral genome. One major question that arose from this observation was whether faithful formation of the strain-specific NP binding profile would impact virus replication. We performed sub-segmental mutational analyses and observed that exchanging nucleotide sequences can alter the NP binding profile (**Figure 2**). Unexpectedly, while NP peaks either ectopically appeared or were ablated at the mutation site, the most striking observation of this study was that alterations in NP binding were not limited to the mutated regions. Instead, NP binding was affected genome-wide on other segments as well despite the fact that their nucleotide sequences remained identical to the wildtype strain (**Figure 3**). We refer to this phenomenon as the ‘butterfly effect of NP packaging’, as minute local changes can produce genome-wide effects (**Tables 1+2**). Finally, we demonstrate that mutant strains, which display a modified NP binding profile and reduced replication fitness, have a defect in segment packaging efficiency and an increase in the formation of semi-infectious particles (**Figures 4–6**).

**Table 2.**
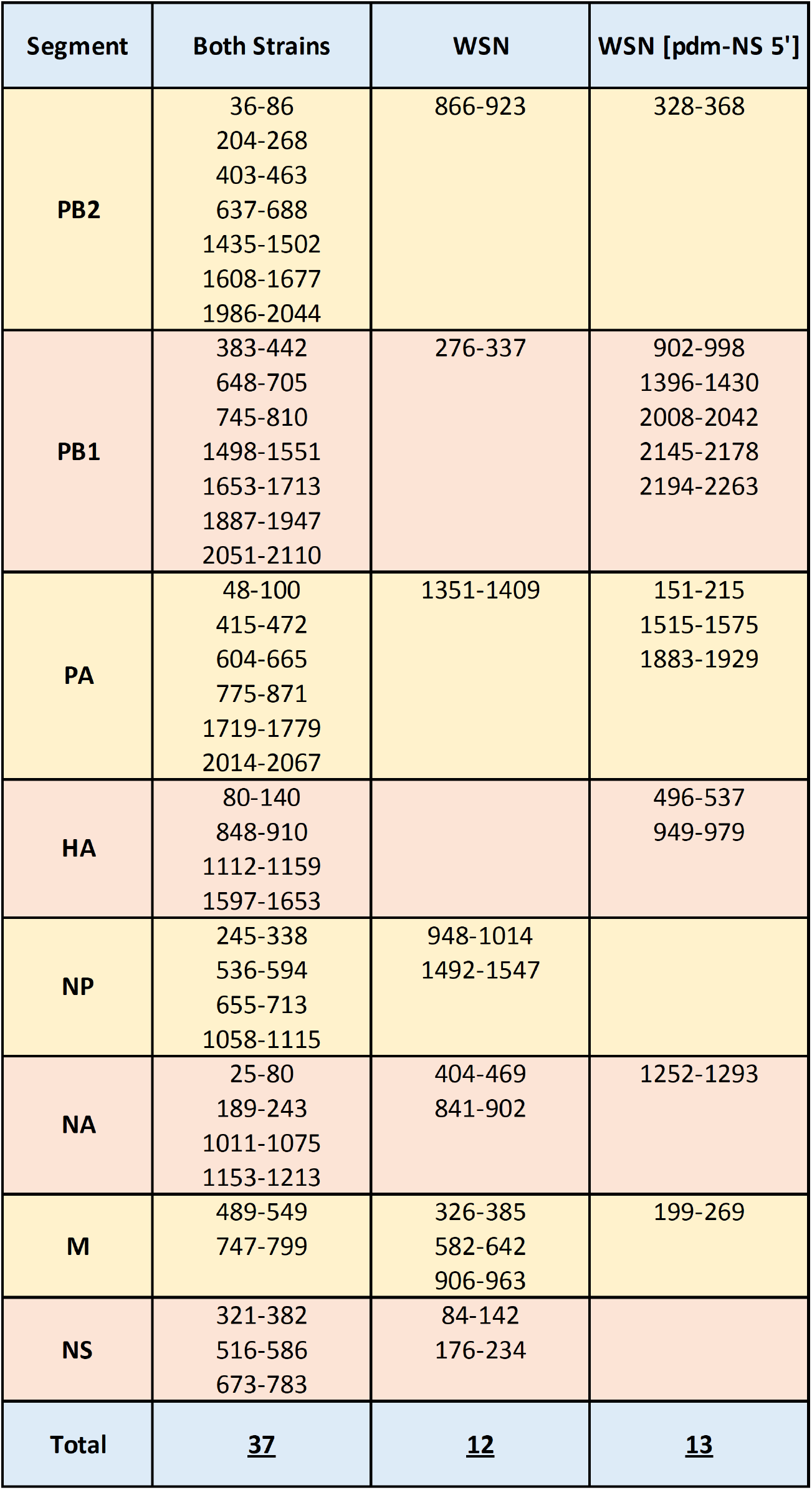
Coordinates of common and unique NP peaks for the wildtype and chimeric NS mutant WSN strains (related to **Figure 3B**).

| Segment | Both Strains | WSN | WSN [pdm-NS 5'] |
| --- | --- | --- | --- |
| **PB2** | 36-86<br>204-268<br>403-463<br>637-688<br>1435-1502<br>1608-1677<br>1986-2044 | 866-923 | 328-368 |
| **PB1** | 383-442<br>648-705<br>745-810<br>1498-1551<br>1653-1713<br>1887-1947<br>2051-2110 | 276-337 | 902-998<br>1396-1430<br>2008-2042<br>2145-2178<br>2194-2263 |
| **PA** | 48-100<br>415-472<br>604-665<br>775-871<br>1719-1779<br>2014-2067 | 1351-1409 | 151-215<br>1515-1575<br>1883-1929 |
| **HA** | 80-140<br>848-910<br>1112-1159<br>1597-1653 |  | 496-537<br>949-979 |
| **NP** | 245-338<br>536-594<br>655-713<br>1058-1115 | 948-1014<br>1492-1547 |  |
| **NA** | 25-80<br>189-243<br>1011-1075<br>1153-1213 | 404-469<br>841-902 | 1252-1293 |
| **M** | 489-549<br>747-799 | 326-385<br>582-642<br>906-963 | 199-269 |
| **NS** | 321-382<br>516-586<br>673-783 | 84-142<br>176-234 |  |
| **Total** | **<u>37</u>** | **<u>12</u>** | **<u>13</u>** |

A revised influenza virus genome architecture has recently been proposed, which suggests that NP binds vRNA in a non-uniform and non-random manner [18, 21], but how this apparent NP specificity is achieved remained unanswered. It was thus unclear whether nucleotide changes in vRNA would alter the NP binding profile, or whether the NP binding profile would remain static due to an as-yet unidentified mechanism that would maintain NP binding at the original positions in the vRNA. While we indeed observed changes in NP binding caused by changes to the underlying nucleotide sequences, we also observed that identical nucleotide sequences can have distinct context-dependent outcomes in terms of NP association. This observation is in line with previous *in vitro* studies that demonstrated that NP interacts with RNA in a sequence-independent manner [22, 23]. Furthermore, this finding suggests that NP binding is not governed by the underlying vRNA sequence alone, but subject to a more complex layer of regulation. To explain the global effect on NP binding caused by local changes, we propose that higher order genome organization may dictate NP binding and speculate that intersegmental interactions may contribute to shaping the genome-wide NP binding profile.

Interestingly, we observed strain-specific differences in the impact of NP binding on replication fitness, where alteration of the WSN NS segment resulted in a virus with decreased replication and packaging efficiency, while H1N1pdm did not. These results are particularly surprising, since the 5′ region of H1N1pdm had low NP binding, which we previously proposed would coordinate vRNA-vRNA interactions. Therefore, we would have expected that the formation of NP peaks at this site would disrupt these RNA interactions and impact replication and packaging of this virus. In contrast, it was the NP peak-containing 5′ region of WSN NS whose ablation disrupted packaging efficiency and viral replication. While counter to our initial hypothesis, these data may provide a more nuanced view of how NP binding may coordinate inter-segmental interactions. In addition, NP-vRNA binding, as examined here, is only a single aspect of the complex coordinative effort that regulates viral replication and packaging. Overall, strain-distinct characteristics may provide the H1N1pdm strain with more flexibility in packaging, so that the pdm [WSN-NS 5′] chimeric virus could gain NP peaks and yet replicate as efficiently as the wildtype H1N1pdm strain. In support for the role of strain background in packaging plasticity, an increased number of alterations were found in the NP-vRNA binding profile for the pdm [WSN-NS 5′] mutant as compared to the WSN chimeric mutant (**Tables 1+2**), which could indicate that the H1N1pdm background is more amenable to modulations of its NP binding profile. Future studies examining the relationship between NP binding profile and intersegmental interactions, using recently developed technologies to study *in vivo* RNA-RNA interactions employing high-throughput sequencing [29], will help elucidate how genome organization affects virus replication.

## Materials and Methods

### Generating mutant virus strains and measuring viral growth curves

Madin-Darby canine kidney (MDCK) cells were maintained in Minimum Essential Medium Eagle (Sigma-Aldrich) supplemented with 10 % fetal bovine serum (FBS, Hyclone), 2 mM L-glutamine (Gibco) and 1 % penicillin/streptomycin (Gibco). HEK293T cells were cultured in DMEM containing 10% FBS, 2 mM L-glutamine and 1 % penicillin/streptomycin. Rescue of recombinant A/WSN/1933 (H1N1) and A/California/07/2009 (H1N1) strains were previously described [3, 30]. Mutations in segments were performed by either conventional site-directed mutagenesis or chemical gene synthesis followed by subcloning of mutant constructs into rescue vectors. In brief, HEK293T cells were transfected with of each of the eight bidirectional plasmids containing each of the wildtype or mutant segments from either A/WSN/1933 (obtained from Richard Webby, St. Jude Children’s Research Hospital) or A/California/07/2009 (obtained from Jesse Bloom, Fred Hutchinson Cancer Research Center) using TransIT-Express (Mirus) according to the manufacturer’s instruction. The HEK293T supernatant was harvested and used to inoculate MDCK cells. The MDCK cell supernatants containing recombinant virus were collected (CP1) and used to generate a virus stock (CP2). Virus propagation for HITS-CLIP was generated from the same CP1 stock.

Multicycle growth curves were performed by infecting with a multiplicity of infection (MOI) of 0.01. Confluent MDCK cells were inoculated in triplicate with each virus and incubated for 1 h at room temperature with shaking, after which the inoculum was replaced with 500 μl of serum-free medium with 1 mg/mL TPCK-treated trypsin (Worthington Biochemical Corporation). Samples were titered by tissue culture infectious dose 50 (TCID50) [31] or by standard plaque assay in MDCK cells. All growth curve measurements were performed in at least two independent biological replicates.

### HITS-CLIP and deep sequencing data analysis

HITS-CLIP experiments were performed as described [18, 32]. In brief, virions in clarified culture medium were irradiated with UV light at 254 nm (400 mJ/cm^2^ and 200 mJ/cm^2^), followed by ultracentrifugation over a 30% sucrose cushion. Virus particles concentrated from 25 mL of culture supernatant were resuspended in 300 μl PXL buffer (1x PBS, 1% NP40, 0.5% deoxycholate, 0.1% SDS), followed by DNase and partial RNase treatment. Immunoprecipitation was performed with anti-NP antibody (mouse monoclonal antibody MAB8251 from Millipore). Subsequent ligation of 5′ and 3′ adapters, RT reaction and first-round PCR amplification step were carried out as described [32]. The first-round PCR products were converted into an Illumina-compatible deep sequencing library using the NEBNext Ultra DNA Library Prep Kit (NEB), and deep sequencing was carried out using Illumina’s NextSeq platform. Data analysis was performed as described [32] using the NovoAlign alignment program and mapping the reads to reference genomes available from the NCBI database. NP peaks were called using the tag2peak.pl script of CLIP Tool Kit [33] with the options “-ss --valley-seeking --valley-depth 0.5 and -minPH” to take into account a minimum threshold based on deep sequencing coverage of the sample (i.e. the total number of mapped nucleotides/length of the genome). NP binding profiles of WSN and H1N1pdm strains were taken from our previous publication [18]. Deep sequencing data generated in this study were deposited in the Sequence Read Archive under accession no. SRP151136. At least two biological replicates of HITS-CLIP were performed for each strain, and the NP binding profiles of all replicates were highly correlative with Pearson correlation coefficients ranging from 0.71 to 0.86. The reproducibility of our HITS-CLIP assay on Influenza virus strains has been described previously [18].

### Segment packaging competition assay (nine-plasmid competitive transfections)

Plasmids encoding PB2, PB1, PA, HA, NP, NA, and M of the WSN strain were transfected with two distinct plasmids encoding NS as indicated. One μg of each plasmid was mixed with TransIT transfection reagent in Opti-MEM medium and transfected into HEK293T cells; 6 h post-transfection the media was replaced with fresh Opti-MEM medium. Virus supernatants were collected at 24 h and 48 h post-transfection and pooled, followed by virus concentration by ultracentrifugation on a 30% sucrose cushion. RNA from virus pellet was isolated using phenol-chloroform extraction. Samples were treated with ezDNAse (ThermoFisher), and SuperScript IV One-Step RT-PCR was performed using a primer pair (5′-GTTGTAAGGCTTGCATAAATG-3′ and 5′-TACAGAGATTCGCTTGGAGA-3′) that anneals to conserved sequences in both NS variants to amplify in an unbiased manner a 193-bp region of the packaged NS segments encompassing the variant region. The amplicons were then converted into an Illumina-compatible library using NEBNext Ultra II DNA Library Prep Kit (NEB) followed by deep sequencing to determine the incorporation ratio of the two NS variants in progeny viruses. 3.0 - 7.1 x 10^5^ sequence reads were analyzed for each experiment.

### Fluorescence *in situ* hybridization of influenza NS and M segments

FISH was performed on infected cells as previously described [3, 28] using probes against M and NS vRNA segments conjugated to Alexa Fluor 488 and Quasar 570, respectively (Biosearch Technologies). Alexa Fluor 488-phalloidin (Life Technologies) was used to mark the plasma membrane. An Olympus FluoView FV1000 confocal microscope with a 60x oil immersion objective was used to acquire stacks of each cell with z intervals of 0.17 μm. Voxel spacing was approximately 50 x 50 x 170 nm to ensure high resolution images for subsequent analysis. All imaging experiments were performed at least twice and a minimum of five representative cells were analyzed.

3D confocal stacks of FISH were background subtracted and deconvolved with Huygens Professional (version 16.05; Scientific Volume Imaging B.V.) at 40 iterations per deconvolution assuming a signal-to-noise ratio of 20. The images were then analyzed using Imaris software (version 8.4.1; Bitplane AG). DAPI marks the cell nucleus, and the signal was used to create a surface and mask the vRNA signal from the nucleus. The ‘Spots’ feature was used to assign a spot for each FISH probe above 2x the mean fluorescence intensity standard deviation provided by Imaris for each channel to provide an unbiased approach. Cell contours were defined manually using the phalloidin staining. Colocalization of M and NS spots were defined using an Imaris Xtension program called “Colocalization of Spots” within 300 nm (the size of our diffraction limit pixel size). The Imaris Cell feature allowed for integration of the cell contour, nuclear surface, and cytoplasmic colocalized and non-colocated spots. This step ensured that only the signal from a given cell was analyzed for colocalization. The statistics were exported and analyzed in PRISM for each cell.

### Hemagglutination assay

A V-bottom 96-well microtiter plate was used to make 2-fold serial dilutions of virus in PBS. An equal volume of 0.5% turkey red blood cells (RBC) was added and incubated for 30 minutes at room temperature. Settling of the RBC to form a button at the bottom of the well was recorded as negative, whereas hemagglutination (RBC staying in suspension) was assigned a positive result. The highest dilution of virus that caused complete hemagglutination was considered as the end-point in HA titration.

### Western blot analysis on virions

Equivalent plaque forming units (PFU) were concentrated by ultracentrifugation on a 30% sucrose cushion. Virus particles were then resuspended in the same volume of NP40 lysis buffer (50 mM Tris pH 7.4, 150 mM NaCl, 0.5 mM EDTA, 0.5% NP40) for Western blot analysis. Membranes were probed with primary antibodies mouse anti-NP (Millipore; MAB8251), mouse anti-Matrix Protein (Kerafast, Inc.; EMS009) or goat anti-Influenza A Virus (Abcam; ab20841) at a dilution of 1:1000. The appropriate HRP-conjugated secondary antibodies (Jackson Laboratories) were used at a dilution of 1:4000. For quantitation, the pixel intensity of each band was determined using the ImageJ software (NIH) and then normalized to the indicated control.

### Relative quantification of viral RNA segments per PFU

vRNA was extracted from virus supernatant containing the same amount of PFU using PureLink Viral RNA/DNA Mini Kit (Invitrogen). vRNA was reverse transcribed with Uni12/13 specific primers using Superscript IV First-Strand Synthesis System (Invitrogen) as per manufacturer’s instructions. The synthesized cDNA was mixed with specific primers for each segment in SYBR Green PCR Master Mix (Applied Biosystems) and the reaction performed on a 7900HT Fast Real-Time PCR System (Applied Biosystems).

## Acknowledgements

S.S.L. and N.L. are supported by the Charles E. Kaufman Foundation. D.J.S and V.S.C are supported by the National Institute of Health [grant number U01AI124303]. We thank Elizabeth McGrady and other members of the Lakdawala lab for technical support and helpful discussions. S.S.L. and N.L. are named inventors on a patent application describing the use of antisense oligonucleotides against specific NP binding sites as therapeutics.

